# A highly diverse, desert-like microbial biocenosis on solar panels in a Mediterranean city

**DOI:** 10.1101/029660

**Authors:** Pedro Dorado-Morales, Cristina Vilanova, Juli Peretó, Franscisco M. Codoñer, Daniel Ramón, Manuel Porcar

## Abstract

Microorganisms colonize a wide range of natural and artificial environments although there are hardly any data on the microbial ecology of one the most widespread man-made extreme structures: solar panels. Here we show that solar panels in a Mediterranean city (Valencia, Spain) harbor a highly diverse microbial community with more than 500 different species per panel, most of which belong to drought-, heat- and radiation-adapted bacterial genera, and sun-irradiation adapted epiphytic fungi. The taxonomic and functional profiles of this microbial community and the characterization of selected culturable bacteria reveal the existence of a diverse mesophilic microbial community on the panels surface. This biocenosis proved to be more similar to the ones inhabiting deserts than to any human or urban microbial ecosystem. This unique microbial community shows different day/night proteomic profiles; it is dominated by reddish pigment- and sphingolipid-producers, and is adapted to withstand circadian cycles of high temperatures, desiccation and solar radiation.

## Introduction

Today, photovoltaic panels cover around 4000 square kilometers, and are forecasted to be the world’s main electricity source by 2050 (http://www.epia.org). Solar panels are unique biotopes characterized by a smooth flat glass or glass-like surface, minimum water retention capacity and maximum sunlight exposure, all of which determine circadian and annual peaks of irradiation, desiccation and heat. Extreme natural habitats such as thermal vents, mountain plateaus or hyper arid deserts are known to host microbial biocenoses adapted to those particular selection pressures (1, 2, 3); and artificial or humanized environments, such as industrial reactors (4), radioactive waste (5) or oil spills (6) are also colonizable by specialized microorganisms. However, despite the popularity and the quick spreading of photovoltaic panels, the microbial communities potentially associated to these human-manufactured devices have not been described to date. Our report documents a complete bioprospection and characterization of the microbial community on photovoltaic panels of a Mediterranean city, using high throughput 16S/18S analysis, metagenomic sequencing, metaproteomics, and culture-based characterization of selected isolates.

## Materials and Methods

### Sampling

Sampling was performed during the summer solstice of 2013 and 2014. Sampling consisted of a simple harvesting procedure, by pouring sterile PBS (sodium phosphate buffer) on the panel and immediately harvesting the liquid by strongly scraping the surface with a modified window cleaner with an autoclaved silicone tube measuring 5 mm in diameter. The resulting suspension was collected by using a sterile plastic pipette and transferred to sterile Falcon tubes, placed on ice and immediately transported to the lab. In 2013, nine samples from solar panels on the three campuses of the University of Valencia (Valencia, Spain) were collected at noon (2 PM) and pooled. Air temperature was 33 C and relative humidity was 60%. In 2014, samples from three independent solar panels in a single location (Faculty of Economics, University of Valencia, Valencia, Spain) were harvested at noon (2 PM) and at night (4 AM) for the proteomic studies, whereas three additional samples (panels 1, 2 and 3) were sampled and used for both 16S/18S rRNA taxonomic identification and metagenomics (Figure 1A). Air temperature and relative humidity were 32 C and 56% (2 PM), and 23 C and 83% (4 AM). The average temperature of the panels surface during the sampling process (at 2 PM) was 51 C. The average solar irradiance in Valencia at 2 PM is 461.3 W/m^2^, whereas the accumulated solar irradiance during an average day is 19.6 kJ/m^2^.

**Figure 1:**
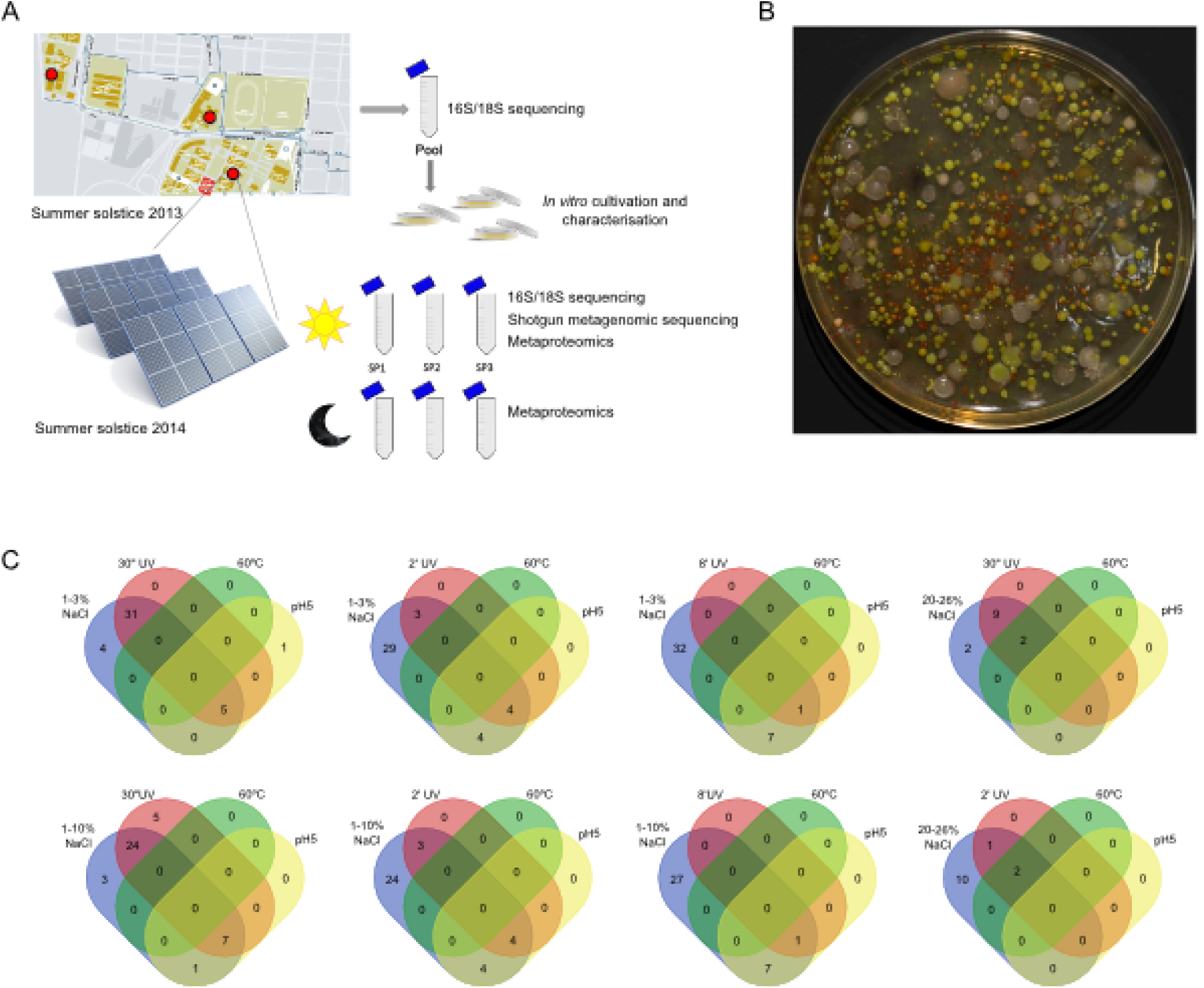
*Characterization of the solar-panel microbiome. Sampling of photovoltaic panels carried out on the three campuses of the University of Valencia in 2013 and 2014, and experimental set up (****A****). Microbial colonies growing on LB incubated at room temperature for two weeks (****B****). Venn diagrams (****C****) displaying the number of isolates exhibiting resistance to heat shock (60C), low pH (5) and different NaCl concentrations (w:vol) and UV pulses (340 W cm-2).*

### Microbiological media and growth conditions

Aliquots of 100 L from the solar-panel samples were spread on Petri dishes containing LB medium (composition in g/L: peptone 10.0, NaCl 10.0, yeast extract 5.0) or marine medium (composition in g/L: peptone 5.0, yeast extract 1.0, ferric citrate 0.1, NaCl 19.45, MgCl_2_ 5.9, Na_2_SO4 3.24, CaCl_2_ 1.8, KCl 0.55, NaHCO_3_ 0.16, KBr 0.08, SrCl_2_ 0.034, H_3_BO_3_ 0.022, Na_4_O_4_Si 0.004, NaF 0.024, NH_4_NO_3_ 0.0016 and Na_2_HPO_4_ 0.008), and incubated at room temperature for 7 days. Individual colonies were independently re-streaked on new media and pure cultures were finally identified through 16S rRNA sequencing and crio-preserved in 20% glycerol (v/v) until required. A total of 53 bacterial strains were characterized under stress conditions and serially confronted with each other in solid medium to detect interactions in terms of resistance to harsh conditions.

### Strains stress tests

Each strain was subjected to a range of stress assays to test tolerance to salinity, heat, low pH, and UV radiation. Overnight liquid cultures were adjusted to an OD600 value of 0.03. Then, stress tests were carried out by plating several 20 L droplets of the diluted culture on LB or marine agar with the following modifications. In the case of salinity stress, increasing amounts of NaCl (from 1 to 9% w/v) were added to the media (final concentration ranging from 1 to 26% w/v). To test pH resistance, culture media was adjusted to pH 5, 6, and 8. In the case of heat, plates were incubated overnight at 60 C; whereas resistance to radiation was tested by applying UV pulses of different length (30 s, 2 min, and 8 min) with a VL-4C lamp (254 nm, 340 W cm^−^2; Labolan, S.L., Spain). The XL1-Blue *E. coli* strain was used as control. For each experiment, two independent replicates were performed. In order to detect inhibition or synergistic effects between strains, experiments were performed as described above and strain suspensions (20 ul) were closely (3 mm) placed on the same dish.

### DNA isolation

Selected DNA purification methods were used to process the solar panels samples and DNA yields were compared (data not shown). Metagenomic DNA was isolated using the Power Soil DNA Isolation kit (MO BIO Laboratories) following the manufacturers instructions with an additional pretreatment with DNA-free lysozyme at 37 C for 10 min. The quantity and quality of the DNA was determined on a 1.5% agarose gel and with a Nanodrop-1000 Spectophotometer (Thermo Scientific, Wilmington, DE).

### PCR amplification and 16S/18S rRNA massive sequencing

A set of primers adapted to massive sequencing for the Ion Torrent platform (Lifetechnologies) were used to capture 16S (modified from (7)) and 18S (modified from (8)) rRNA from the solar-panel DNA extraction in a PCR reaction. PCR reactions were performed with 30 ng of metagenomic DNA, 200 M of each of the four deoxynucleoside triphosphates, 400 nM of each primer, 2.5 U of FastStart HiFi Polymerase, and the appropriate buffer with MgCl2 supplied by the manufacturer (Roche, Mannheim, Germany), 4% of 20 g/mL BSA (Sigma, Dorset, United Kingdom), and 0.5 M Betaine (Sigma). Thermal cycling consisted of initial denaturation at 94C for 2 minutes followed by 35 cycles of denaturation at 94C for 20 seconds, annealing at 50C for 30 seconds, and extension at 72C for 5 minutes. Amplicons were combined in a single tube in equimolar concentrations. The pooled amplicon mixture was purified twice (AMPure XP kit, Agencourt, Takeley, United Kingdom) and the cleaned pool requantified using the PicoGreen assay (Quant-iT, PicoGreen DNA assay, Invitrogen). Subsequently, sequencing on the Ion Torrent platform was performed at LifeSequencing S.L. (Valencia, Spain).

### Shotgun metagenomic sequencing

The metagenomic DNA of two of the solar panels sampled in 2014 (solar panels 1 and 3, from which enough DNA was available) was shotgun sequenced. A Nextera Illumina library was built from 100 ng total DNA following the protocol indications marked by Illumina. Those libraries were sequenced in a MiSeq sequencer (Illumina) at Lifesequencing SL, in a combination of 500 cycles, in order to obtain 250 bp paired-end sequences.

### Bioinformatic analysis

#### 16S/18S rRNA profiles

The resulting sequences from the taxonomical identification, based on PCR capturing of the 16S and 18S rRNA, were split taking into account the barcode introduced during the PCR reaction, providing a single FASTQ file for each of the samples. We performed quality filtering (Q20) using fastx tool kit version 0.013, primer (16S and 18S rRNA primers) trimming using cutadapt version 1.4.1 and length (minimum 300 bp read length) trimming using in-house perl scripting over those FASTQ files to obtain a FASTQ file with clean data. Those clean FASTQ files were converted to FASTA files and UCHIME (9) program version 7.0.1001 was used to remove chimeras arising during the amplification and sequencing step. Those clean FASTA files were BLAST against NCBI 16S rRNA and fungi database using blastn version 2.2.29+. The resulting XML file were processed using a pipeline developed by Life-sequencing S.L. (Paterna, Valencia, Spain) in order to annotate each sequence at different phylogenetic levels (Phylum, Family, Genera and Species). Statistical analysis was performed using R version 3.1.1. A summary of sequencing statistics and results is available in Table S2.

#### Taxonomic and functional analysis of metagenomic sequences

Two FASTQ files per sample were obtained during the sequencing step, each coming from each of the directions on the paired-end sequencing. Those files were trimmed for adapters and low quality reads using cutadapt version 1.4.1 with the paired-end option. Trimmed sequences were used for taxonomical identification using a local alignment tool against nt database from NCBI as described before (10). The trimmed sequences from each solar panel were also assembled using different combinations of k-mers in Abyss version 1.5.2 (11) and Velvet version 1.2.1 (12) in order to find the best combination. The best assembly in each solar panel was used to perform a prediction of ORFs by using MetaGeneMark (13). BLASTP against nr NCBI database was used for annotation and webMGA (14) for COG assignation. All our data have been deposited in the MG-RAST server, and is publicly available under accession numbers 4629146.3 and 4629747.3. In order to compare the taxonomic profile of solar panels with other environments, the taxonomic information of 25 metagenomes belonging to different habitats was obtained from the MG-RAST server (IDs 4455835.3, 4455836.3, 4477803.3, 4477872.3, 4441205.3, 4445129.3, 4445126.3, 4477903.3, 4477901.3, 4514299.3, 4543019.3, 4543020.3, 4441363.3, 4441215.3, 4441214.3, 4441679.3, 4447192.3, 4447102.3, 4497390.3, 4497389.3, 4516651.3, and 4516403.3). The different profiles were processed with MEGAN. Data were normalized, and the distances between pairs of profiles calculated with the Bray-Curtis method. Finally, the calculated distances were used to build a Principal Coordinates Analysis. We employed the Statistical Analysis of Metagenomic Profiles (STAMP) (version 1.08; Faculty of Computer Science, Dalhousie University) software to compare the functional profile (according to subsystems categories) of our samples with those of the metagenomes previously cited. The functional data of metagenomes 4455835.3, 4455836.3, 4477803.3, 4477873.3, 4477903.3, 4477904.3, 4477901.3, 4514299.3 was poor or absent, and was thus eliminated from the analysis. This comparison was represented in a heatmap, where the different metagenomes are clustered according to their similarity. The functional contents of solar panels 1 and 3 were compared to each other with a Fishers exact test combined with the Newcombe-Wilson method for calculating confidence intervals (nominal coverage of 95%). As a multiple-hypothesis test correction, a false-discovery-rate (FDR) method was applied.

#### Pangenome reconstruction

Trimmed sequences from both solar panels for the total DNA experiment were blasted against a database containing all sequences for the genera Thermo/Deinococcus. Only sequences with a positive hit in against this database were used for assembly them using different k-mers with the Abyss assembler. We performed a genome annotation in different steps i) ORFs were predicted with GeneMark version 3.25 (13), ii) rRNA version 1.2 (15) for rRNA prediction, and iii) tRNA-Scan (16) for tRNA prediction. The functional annotation using COG classification was performed using webMGA. DNAPlotter (17) from the Artemis Package was used to represent a circular map of the pangenome.

### Proteomics

Protein samples were precipitated with TCA (trichloroacetic acid) and pellets were dissolved with 75 L of 50 mM ABC (ammonium bicarbonate). The protein concentration in the samples was determined by fluorometric analysis. Then, 10 g of each sample were digested as described in the following protocol. Cysteine residues were reduced by 2 mM DTT (DLDithiothreitol) in 50 mM ABC at 60C for 20 min. Sulfhydryl groups were alkylated with 5 mM IAM (iodoacetamide) in 50 mM ABC in the dark at room temperature for 30 min. IAM excess was neutralized with 10 mM DTT in 50 mM ABC, 30 min at room temperatura. Ecah sample was subjected to trypsin digestion with 250 ng (100 ng/l) of sequencing grade modified trypsin (Promega) in 50 mM ABC at 37C overnight. The reaction was stopped with TFA (trifluoroacetic acid) at a final concentration of 0.1%. Final peptide mixture was concentrated in a speed vacuum and resuspended in 30L of 2% ACN, 0.1% TFA. Finally, 5 l of each sample were loaded onto a trap column (NanoLC Column, 3 C18CL, 75um x15cm; Eksigen) and desalted with 0.1% TFA at 2l/min during 10 min.

The peptides were then loaded onto an analytical column (LC Column, 3 C18CL, 75um×25cm, Eksigen) equilibrated in 5% acetonitrile 0.1% FA (formic acid). Elution was carried out with a linear gradient of 5:35% B in A for 40 min (A: 0.1% FA; B: ACN, 0.1% FA) at a flow rate of 300 nl/min in a label free mode. Peptides were analyzed in a mass spectrometer nanoESI qQTOF (5600 TripleTOF, ABSCIEX). The tripleTOF was operated in informationdependent acquisition mode, in which a 0.25s TOF MS scan from 350 to 1250 m/z, was performed, followed by 0.05s product ion scans from 100 to 1500 m/z on the 25 most intense 25 charged ions.

ProteinPilot default parameters were used to generate a peak list directly from 5600 TripleTof wiff files. The Paragon algorithm of ProteinPilot was used to search NCBI protein database with the following parameters: trypsin specificity, cysalkylation, no taxonomy restriction, and the search effort set to through. To avoid using the same spectral evidence in more than one protein, the identified proteins were grouped based on MS/MS spectra by the ProteinPilot Progroup algorithm. The PeakView v 1.1 (ABsciex) software was used to generate the peptides areas from Protein Pilot result files and to perform a principal component (PCA) and a ttest analysis.

## Results

Culturing of solar panel samples from the 2013 solstice on both rich (LB) and marine media yielded a relatively high number of colony forming microorganisms, mostly bacteria, displaying a wide range of color and shapes. Many of the isolates displayed red, orange or pink pigmentation (Figure 1B), particularly those incubated on marine agar. A total of 53 pigmented isolates were selected and subjected to taxonomic (16S rRNA) characterization and tested for resistance to heat (incubation at 60C), UV exposure (2 to 30 s pulses of a 340 W cm^−^2 UV light), high NaCl contents (1 to 26%) and different pH values (5 to 9).

Figure 1C shows that, in general, the solar-panel isolates displayed strong resistance to very high salt concentrations, moderately high resistance to low pH and relatively low resistance to UV light or extreme (60C) heat. Interestingly, during these characterization assays we were able to identify isolates able to restore the growth of nearby isolates under conditions of extreme salt or pH values, in the latter case because of local buffering of the pH of the plate (Figure S1). Full characterization of the 53 isolates is provided in Table S1 and Figure S2.

The taxonomic composition of bacterial and eukaryotic taxa was first studied through 16S and 18S rRNA genes massive sequencing; the results are shown in Figure 2. As many as 800 different bacterial species were identified in the 2013 pool (nine solar panels from different locations within the University of Valencia); and around 500 different species were found in each of the individual panels sampled from a single building in 2014 (Table S2). Two orders, Sphingobacteriales (families Flexibacteriaceae and Sphingomonadaceae) and Deinococcales comprised the highest number of species. *Deinococcus, Sphingomonas, Novosphingobium* or *Hymenobacter* were the dominant genera, with 9.2% or 28% of the assigned sequences (*H. chitinivorans* in panel 1 and *D. hopiensis* in panel 3, respectively). The remaining sequences were distributed among 17 phyla and 146 families. Other well-represented genera, in order of abundance, were *Rubellimicrobium, Adhaeribacter, Acidicaldus, Segetibacter,* or *Modestobacter.* In the case of fungi, lower biodiversity was found (Fig. 2B). Taxonomic eukaryotic profiles were dominated by the phylum Ascomycota and the families Pleosporaceae and Teratosphaeriaceae, with genera *Phaeothecoidea* and *Alternaria* representing the majority in the 2013 and 2014 samples, respectively.

**Figure 2:**
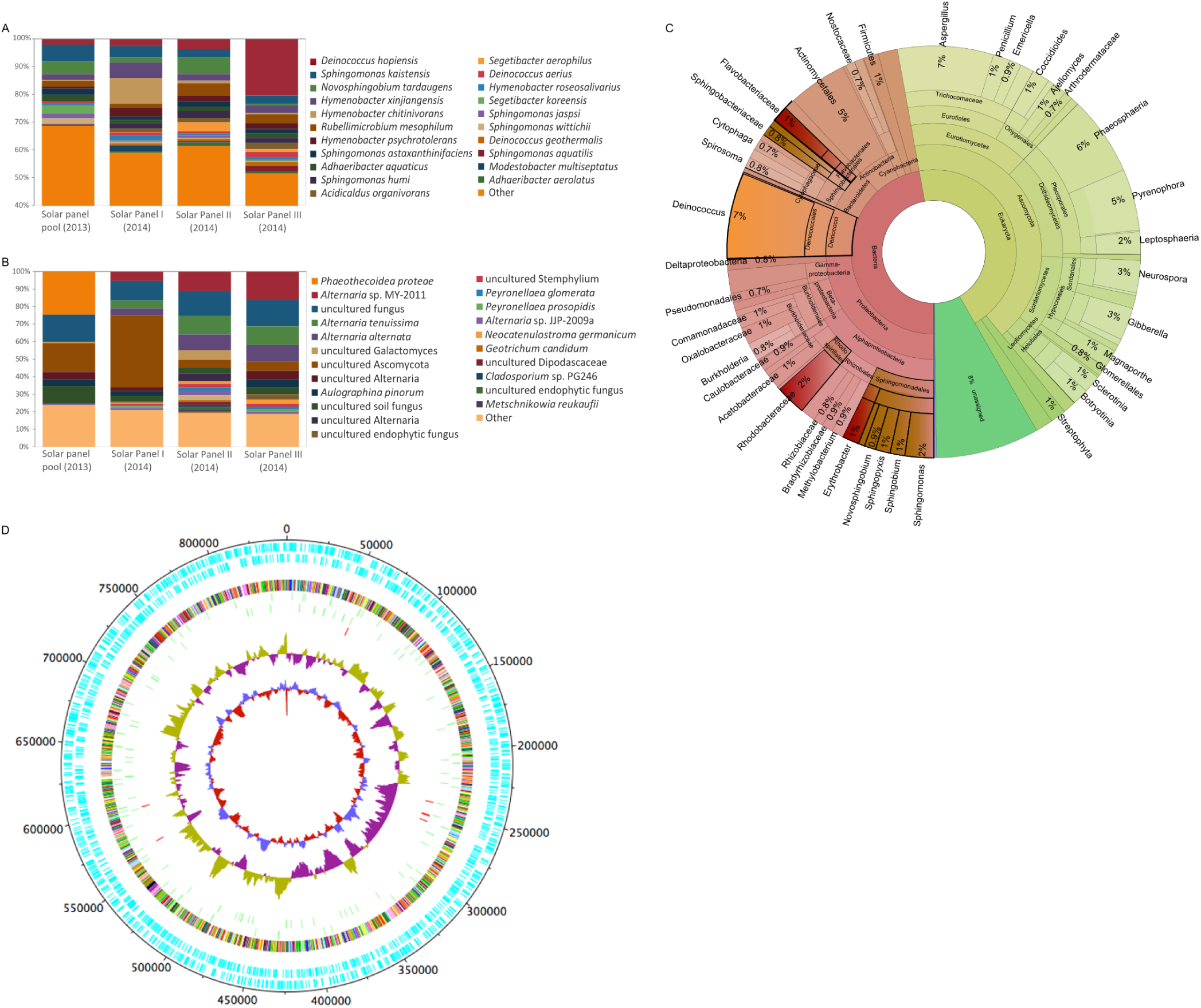
*Taxonomic diversity of solar-panel biocenosis, deduced by culture-independent techniques. Diversity of bacteria (****A****) and fungi (****B****) analyzed by 16S and 18S rRNA gene sequencing, respectively, of solar panels sampled during the Summer solstice of 2013 and 2014. Histograms show the relative abundance (%) of the species identified, as described in materials and methods. Species representing less than 1% of total reads were clustered and labeled as ”Other”. Taxonomic diversity of one of the panels (panel 3) sampled in the summer solstice of 2014 as deduced from shotgun metagenomic sequencing (****C****). Carotenoid and sphingolipid-producing bacteria are highlighted in red and brown, respectively. Radiation-resistant phyla are highlighted in orange. The taxonomic diversity of solar panel 1 is represented in Figure S3. Circular representation of the Deinococcus solar panel pangenome (****D****) obtained from the metagenomic sequences of panels 1 and 3. The map includes (from the outer to the inner circle) the ORFs in forward and reverse sense, a colour-coded COG functional annotation, the predicted tRNAs and rRNAs, the GC count, and the GC skew.*

In 2014, the metagenomic DNA of two independent solar panels was shotgun sequenced. As shown in Figure 2C, the bacteria:fungi ratio was close to 50%, and species distribution was similar to that found for 16S and 18S sequencing. A summary of sequencing statistics and diversity indexes can be accessed in Table S3. Genus *Deinococcus,* one of the clearest taxonomic markers of extremophily, again proved highly abundant in all our samples. We analyzed Deinococcus sequences from our metagenomic analysis and a draft *Deinococcus* solar panel pangenome of 2098 contigs was obtained, covering more than 0, 8 Mb (25%) of standard *Deninococcus* genomes (3.3Mb with 2 chromosomes and 2 plasmids), with 2166 and 149 ORFs and tRNAs, respectively (Figure 2D). The low identity level of the solar panel pangenome with previously sequenced *Deinococcus* species strongly suggests that at least one previously undescribed Deinococcus species is present in the sampled panels.

Regarding the functional profile, that of two independent solar panels (1 and 3) was deduced from the metagenomic data, and statistically analyzed with the STAMP software. When compared to a range of metagenomes from diverse habitats, solar-panel functional profiles clustered together with those described for polar microbial mat and saline desert datasets, and distant to other environments such as air or sediments (Figure 3A). Both solar panels proved very similar to each other in terms of functions, as shown in Figure 3B. The bioactivity of the biocenosis was studied through a metaproteomic analysis conducted on solar panels sampled at noon (solar time) and at night (4 AM). Protein composition differed between the day and night samples (Figure 3C). Significantly, a protein involved in modulating bacterial growth on surfaces and biofilm formation (18) (diguanylate cyclase) was particularly abundant. Also among the more expressed proteins, we identified fungal and bacterial enzymes involved in respiration and ATP synthesis or ribosomal proteins (bacterial L7/L12 and archaeal L7, the latter being a moonlighting protein involved in rRNA processing (19). Other abundant proteins have been reported to confer resistance/tolerance to the extreme conditions found in solar panels, namely, salt stress and drought (membrane-bound proton-translocating pyrophosphatase mPP) (20), nutrient starvation (mPP and cold-shock protein) (21), heat-shock (molecular chaperone GroEL), as well as proteins involved in the preservation of membrane integrity under harsh conditions (22, 23) (S-layer protein, lipoprotein 1; Table S4).

**Figure 3:**
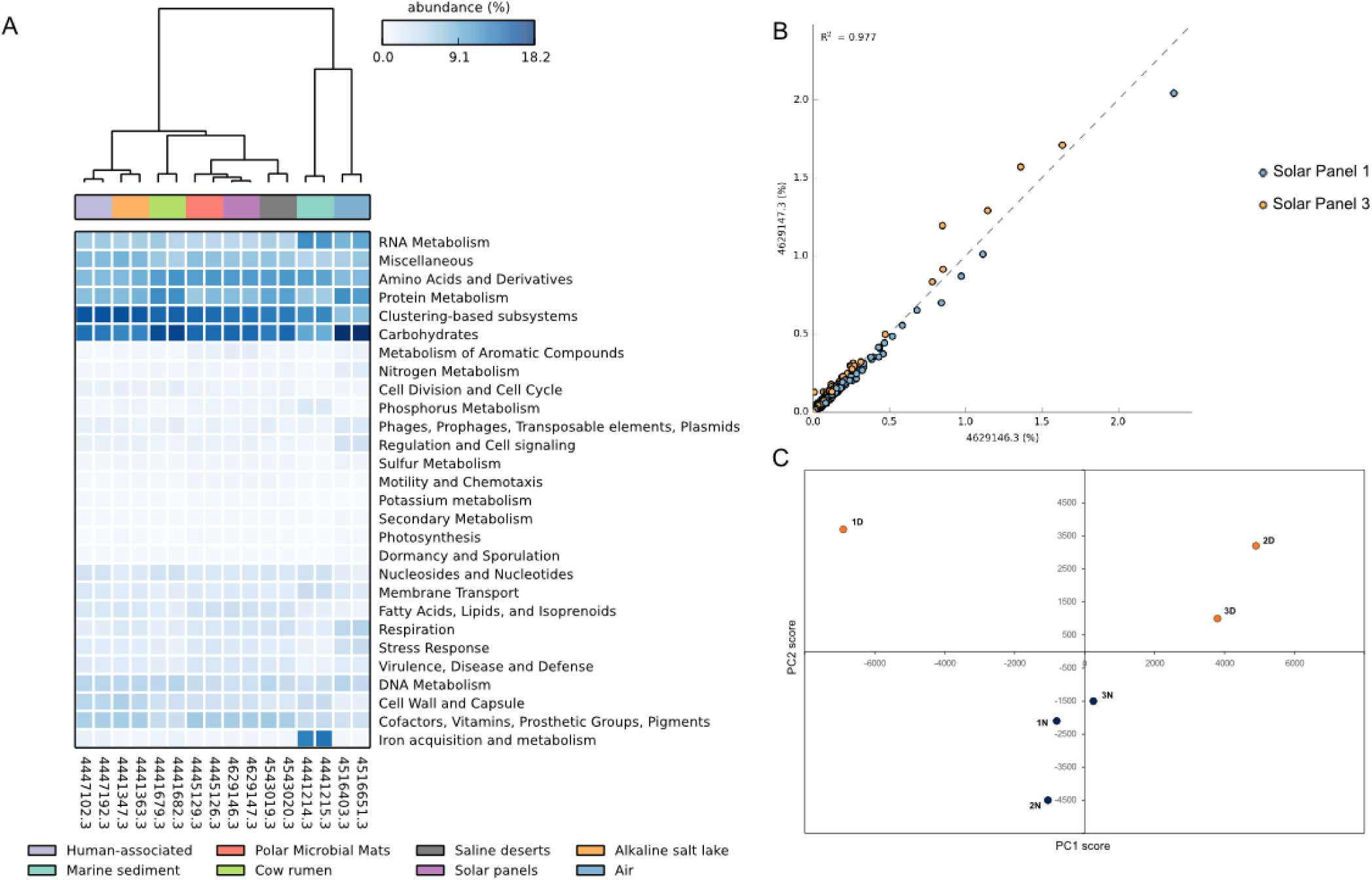
*Functional analysis of the solar-panel metagenomes. Heatmap representation (****A****) based on the functional profiles of solar panels 1 and 3 compared with a range of metagenomes from different environments. Set of functions in the metagenomes of solar panels 1 and 3 (****B****). Each dot corresponds to one function of the subsystems classification. Principal Component Analysis (****C****) performed with the proteomic profile of solar panels sampled at noon (yellow dots) and night (dark blue).*

## Discussion

Despite the harsh conditions to which microorganisms deposited -or permanently inhabiting-the solar panels of a Mediterranean city during summer, standard culturing resulting in important microbial growth, with a diversity in shape, color and textures of colony forming microorganisms from the 2013 solstice suggesting high biodiversity of the environment. Although culturable isolates are typically only a small fraction of the global biocenosis, we were able to identify several strain-to-strain effects that proved able to restore the sensitivity of neighboring isolates to stress factors (salinity and low pH). These results suggest that microbial interactions and the particular physical location of microorganisms on the solar panels, rather than individual cell properties, might play a major role in bacterial survival on solar panels.

High throughput sequencing allowed confirming the high diversity of the habitat in the form of a sun-adapted taxonomic profile. Indeed, and in accordance with the majoritary phenotype observed within culturable isolates, most of the species identified by high throughput sequencing, and particularly the most frequent ones, are known to produce pink (*H. xingiangensis* (24)), *H. psychrotolerans* (25)), orange (*Sphingomonas humi* (26)), orange-red (*S. kaistensis* (27)) or reddish pigments (*Hymenobacter chitinivorans* (28), *Rubellimicrobium mesophilum* (29)), in most cases carotenoids; as well as sphingolipids (*Sphingomonas* spp. (26, 27), *Novosphingobium* spp. (30)). Carotenoids have been reported to play a major role in radiation tolerance in bacteria (31) and sphingolipids have recently been described to mediate bacteria-to-silica and polyamide adhesion (32). Therefore, carotenoids and sphingolipids are candidates accounting for the sunlight resistance and fixation properties that microorganisms need to survive on a smooth, south-facing surface.

A review of the ecology of the main bacterial taxa we identified gives more insights of the extremophile character of the solar panel bacteriome. Indeed, several of the most frequent *Deinococcus* spp. and other solar-panel bacteria have been described as inhabitants of relatively mild desertic areas as well as polar environments. *D. hopiensis* was isolated from the Sonora desert (33), while other *Deinococcus* species that we detected were first described in the Sahara desert (34). Regarding other very abundant species, such as *Hymenobacter xingiangensis* or *Sphingomonas kaistensis,* they have previously been reported on the high Tibet plateau (25), on dry Antarctic valleys (35), or in the Chinese desert of Xingiang (24). Others were first reported in high salinity areas (36), thermal springs (37) or during bioprospections of soil (38, 39), or air samples (40, 41). A systematic review of the locations where the 50 most abundant solar panel bacteria were first isolated reveals their adaptation to extreme environments: most of them occur in drought, high radiation and/or high temperature habitats. Most of the species we found on solar panels were originally reported to inhabit a relatively narrow geographical band in the temperate zone of the Northern hemisphere. The distribution of others in the dry Antarctic valleys suggests a major role of radiation as a key selective factor (Figure 4A), and the PCoA analysis of the solar-panel taxonomic profile compared with other metagenomes reveals a clear link with extremophile environments, such as temperate and cold deserts (Figure 4B).

**Figure 4:**
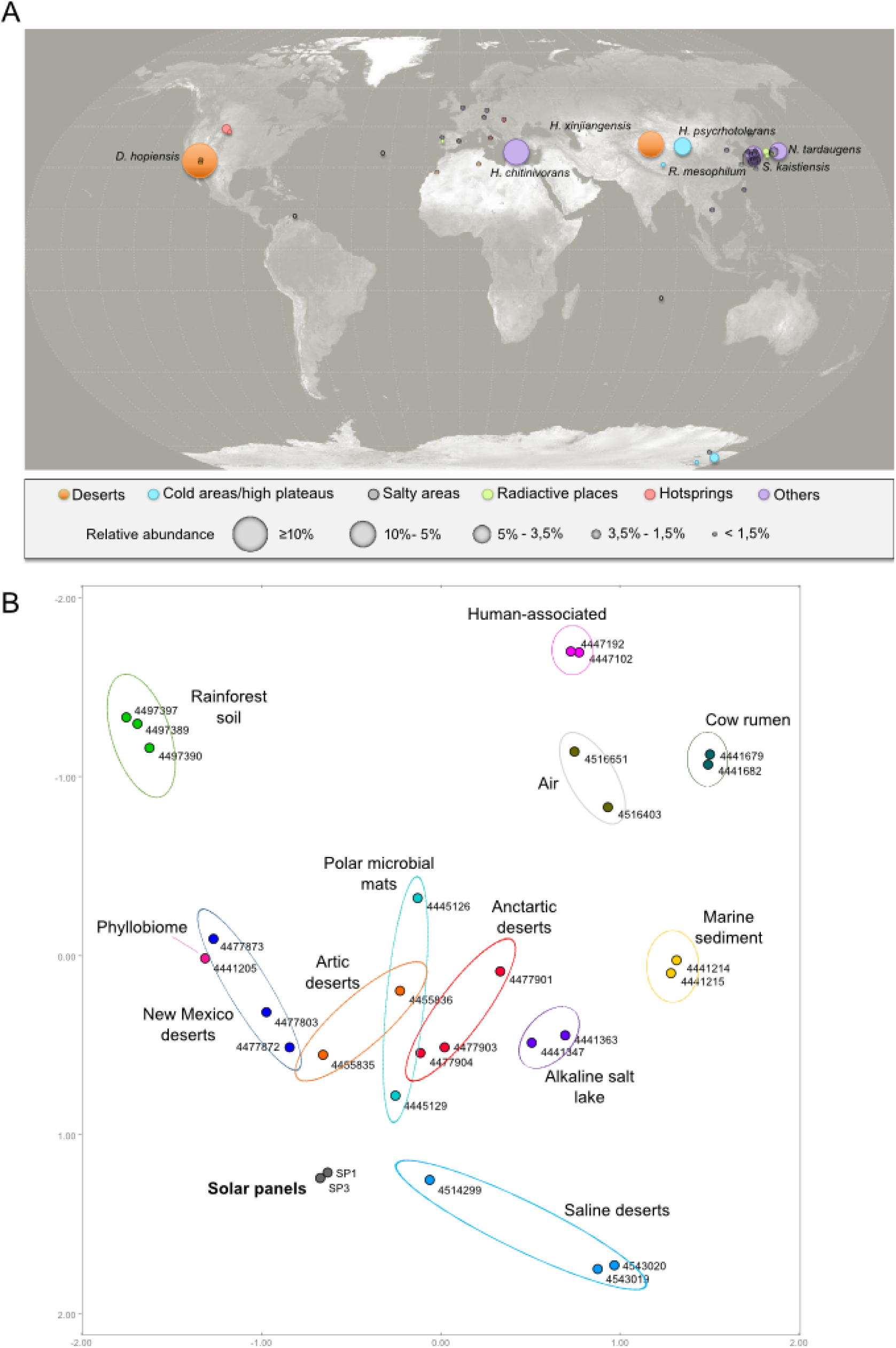
*Biogeographical context of the solar-panel microbiomes as deduced from their taxonomic profile. Geographic distribution (****A****) of the 50 most abundant (more than 1% of the reads) bacterial species detected by high-throughput sequencing of 16S rRNA amplicons in solar panels sampled, at the University of Valencia in 2014. Each circle corresponds to a different species and the size of the circles is proportional to the number of reads. Species found at a frequency higher than 3.5% are shown. Colors indicate type of environment. Principal Coordinates Analysis (****B****) performed with the taxonomic profile of a range of metagenomes from diverse ecosystems. The solar-panel metagenomes (panels 1 and 3 from the 2014 sampling, grey dots) map within desert and circumpolar metagenomes.*

Regarding fungi, the ecological niches of several of the most frequent genera (*Neocatenulostroma, Xenophacidiella* and *Metschnikowia*) are sunny habitats, such as the phylloplane: (42, 43); or the surface of rocks (*Coniosporium* spp., particularly abundant in the 2013 samples) (44). As in the case of bacteria, this taxonomic profile strongly suggests sunlight exerts a major selective pressure, shaping the fungal community on the panels. The abundance and diversity of microorganisms in solar panels can be solely due to wind-deposition or correspond to an in situ active ecological community. Protein composition differed significantly between the day and night samples (Figure 3C), implying that the microbial communities populating the solar panel surface are biologically active. The abundance of proteins involved in resistance to harsh conditions and biofilm formation on surfaces proves the presence of stress-response mechanisms in the microbial communities inhabiting solar panels. These results, along with the abundance of radiation-resistant taxa and the desert-like taxonomic profile of the solar samples, that strikingly plot within desert microbiomes (Figure 4B), strongly suggest an in situ adaptation from (probably) wind-transported microorganisms that are immediately subjected to selection with radiation, heat and dessication as main shapers of this microbial ecosystem. Indeed, the solar panels microbiome proved taxonomically very distant from that associated to air samples from a similar latitude (see metagenomes 4516651.3 and 4516403.3 in Figure 4B). Furthermore, a recent analysis of air samples in Sardinia (45), a known Mediterranean crossroad of dust-conveying winds from Saharan Africa, revealed an extremely low abundance of extremophiles such as Deinococcus-Thermus species (less than 1% of relative abundance, in contrast to up to 30% in solar panels). Taken together, all these results strongly suggest that the diverse biocenosis on solar panels we report here is not a mere consequence of physical accumulation of air-driven microorganisms, but a resident microbial community adapted to desert-like selection pressures.

This is the first report of a highly diverse microbial community on solar panels. A recently published study reported limited microbial diversity on solar panels in Brazil, including some fungal species which hindered the panels photovoltaic efficiency (46). Our data show for the first time that solar panels of a temperate Mediterranean city support a highly diverse and active ecological community, one of the richest extremophile biocenoses described to date. Moreover, this community is metabolically active and displays striking taxonomic and functional similarities with highly irradiated environments: temperate deserts and polar environments. The detailed analysis of the habitats where the solar panel microorganisms have previously been detected indicates their strong adaptation to sun exposure, which can only be partially reproduced by stress characterization on pure microbial cultures. Microbial interactions (including pH and salinity tolerance restoration), physical effects such as shading, DNA repair mechanisms and production of pigments and adhesion molecules might play a major role in the adaptation of a unique microbial ecosystem to the abrupt circadian cycles in desert-like conditions. This previously undescribed ecosystem is the first urban microdesert reported to date, and it may provide a valuable new source of compounds with biotechnological applications.

## Acknowledgments

Data reported in this paper have been deposited in the MG-RAST server, and are publicly available under accession numbers 4629146.3 and 4629147.3. The authors are indebted to the Vicerrectorat dIn-fraestructures (Universitat de València) for their support before and during sampling. We also thank the proteomics service (SCSIE) of the University of Valencia for its technical support and Fabiola Barraclough for English revision. P.D. and C.V. contributed equally to this work.

